# A model that predicts resistant starch in a dog kibble using a small-scale twin-screw extruder

**DOI:** 10.1101/2021.10.22.465457

**Authors:** Isabella Corsato Alvarenga, Christopher Waldy, Lewis C. Keller, Charles G. Aldrich

**Author notes:** Corresponding author; Kansas State University 003 Waters Hall, Manhattan KS 66506.

## Abstract

The objective of this work was to modify extrusion parameters to yield greater resistant starch (RS) in a kibble and create a model to predict its concentration. A dog food was extruded through a small-scale twin-screw extruder as a central composite design with 6 central points (replicates) and 14 single replicates. There were three factors tested at three levels: corn particle size, extruder shaft speed, and in-barrel moisture (IBM). The remaining processing inputs were kept constant. Chemical and physical starch analyses were performed. A model to predict RS was created using the REG procedure from SAS. Pearson correlations between extrusion parameters and starch analyses were conducted. A model to predict RS was created (R^2^_adj_= 0.834; P < .0001). Both SME and extrudate temperature had a high negative correlation with RS and RVA raw starch. Results suggest that low mechanical energy and high IBM increase kibble RS.

## 1. Introduction

Extrusion is the most common process through which pet foods are produced. It can be classified as a medium shear (screw speed above 100 rpm), medium temperature (55 to 145 ºC), and medium moisture (15 to 30%) process (1). Before cooking, cereals and other dietary ingredients are ground and mixed, then fed to a preconditioner (PC) where water and steam are added and mixed with the dry recipe. This step hydrates the mix and starch granules begin to swell. From the PC the mix is fed into the extruder barrel, the primary cooking apparatus. At this stage, pressure increases, screw rotation transfers mechanical energy to the dough, and additional steam may be added. The moisture content and energy transfer enable the dough to transition from a glassy to rubbery state which traps water droplets within the melt. Water droplets expand as these vaporize when the material exits the extruder and the product goes from high pressure to atmospheric pressure (2). Post extrusion, kibbles are dried to less than 10% moisture and coated with fat and flavors.

The extrusion process conditions such as water and steam additions combined with mechanical energy promote starch gelatinization (3). Past studies have demonstrated that decreasing the amount of mechanical energy (4–6), increasing process water content addition (3), and increasing the starch ingredient particle size (4,5) minimize starch gelatinization, which yields some resistant starches (RS). Resistant starches represent the starch fraction that escapes small intestinal digestion and undergoes fermentation by saccharolytic bacteria within the colon to produce short chain fatty acids (SCFA). Previous data have shown an increased butyrate production from consumption of RS from low shear extrusion (4–6). It is common in the human food industry to produce RS through chemical or enzymatic modification of raw starches, and their effects on health have been extensively studied (7). In pet foods, some studies have explored the health benefits of supplementing RS in dog diets across various breeds (8,9) and a few have attempted to retain RS in extruded foods (4–6). The latter represents an opportunity in the pet food industry. Modifying the process to retain some of the native crystalline resistant starch (type II) and(or) develop retrograded starch (type III) may be more cost effective, as well as provide a health attribute to the food. Therefore, the objectives of the present study were: 1. maximize RS of corn in the final kibble by controlling three processing factors including particle size, water input at the extruder barrel, and extruder shaft speed, 2. create a response surface model that predicts RS concentration based on process inputs, and 3. correlate physical analyses of kibble factors with chemical methods for starch conversion.

## 2. Materials and methods

### 2.1 Diet

A single diet was formulated (Concept5©; CFC Tech Services Inc., Pierz, MN, U.S.A.) to meet the nutrient requirements for adult dogs at maintenance (10). Only the dry ground ingredients were included in the dry mix, and any flavors, fats or oils from the coating step were excluded (Table 1). This was chosen because the focus of the present work was not to feed diets to dogs, but to assess the effects of extrusion processing on starch transformation. Whole yellow corn (Cargill, KS, U.S.A.) was the sole starch ingredient and no fiber ingredients were added to the formula (Table 1). The remainder of the dry mix included chicken meal (Tyson, AR U.S.A.), amino acids, minerals and vitamins (DSM Additive Mfg., Overland Park, KS, U.S.A.).

**Table 1.** Diet formula with ingredients added to the dry mix (as-is), which was fed to the extruder.

| **Ingredient** | **%** |
| --- | --- |
| Whole yellow corn | 75.84 |
| Chicken meal | 23.17 |
| Potassium chloride | 0.46 |
| Vitamin premix | 0.12 |
| Lysine, hydrochloride | 0.12 |
| Sodium chloride, iodized | 0.12 |
| Taurine | 0.06 |
| Mineral premix | 0.12 |

### 2.2 Particle size

Particle analysis of the corn and dry mix was determined by a Morphologi G3 instrument (Malvern Panalytical; Malvern, United Kingdom) using 5 x magnification, 5 mm^3^ sample size and one bar of air pressure for dispersion of the sample.

Particle size distribution of the dry mix and corn samples were determined by rotating tapping sieve analyses (Ro-tap) (11). This procedure involved 100 g sample and rotating-tapping time of 10 minutes. There were 14 sieves in the stack with screen sizes of 3,360, 2,380, 1,680, 1,191, 841, 594, 420, 297, 212, 150, 103, 73, 53 and 37 μm.

### 2.3 Extrusion processing

Extrusion was conducted in a small-scale intermeshing co-rotating twin-screw extruder (Evolum 25; Clextral, Firminy, France) set up as a 24 length to diameter screw design with twin 25 mm-diameter screws (S. Fig 1). This system utilized a 25 L dual cylinder preconditioner with steam and water injection. The rationale for using this small-scale extruder was to optimize data collection with a small amount of raw material across a wide range of settings that would correspond to a larger scale extruder.

The experiment was conducted as a non-rotatable central composite design with 6 replicated center points and 14 single replicates, totaling 20 samples (Johnson and Milliken, 1989; S. Table 1.). The factors were tested at 3 levels (low, medium, high) on the same recipe at three mill screen sizes (0.793 mm, 1.19 mm and 1.586 mm), three pre-conditioner moisture contents (target 20%, 25% and 30% water added to the dry mix) at constant PC feed rate and three screw speeds (400, 800 and 1200 rpm).

The 6 central point replicates were run sequentially (sample 1 to 6; S. Table 1) to facilitate processing. Each time replicates were switched, extruder shaft speed was either increased to 1200 rpm, or decreased to 400 rpm for 5 minutes, then changed back to 800 rpm. This led to some variation in the process as would be expected for replicates not run sequentially. Samples were collected after 5 minutes of setting changes to allow for leveling to steady state condition for moisture and extrudate temperature inside the die. The other single replicates were run in a random order (S. Table 1).

### 2.4 Processing details and data collection

Corn was ground through a hammermill at three screen sizes: 0.793 mm, 1.19 mm and 1.586 mm in order to produce a fine, medium and coarse grind size. In sequence, ground corn was mixed with the other ingredients of the “dry mix” (Table 1) using a 68 kg capacity ribbon mixer (Model 9, Wenger Mfg., Sabetha, KS), and then the complete dry mix was ground again in a hammermill (Jacobson 120-D portable hammermill; Carter Day International Inc., Minneapolis, MN) with a 1.586 mm screen. The grinding of the dry mix was done to assure no large pieces would interfere with the dough flowing through the extruder die.

Extruder data for input and output variables were recorded every 5 minutes (twice) during each sample production and then averaged (Table 3). The dry mix was delivered to the PC at a constant feed rate of 30 kg/h with a PC shaft speed set to 100 rpm during the whole experiment. The PC steam flow rate was set to 4.5 kg/h, while PC water was varied from 1.5 to 4.5 kg/h to achieve the target moisture by the operator. Extruder water was also kept constant at 2 kg/h in all treatments and no steam was added to the extruder. At the end of the extruder barrel knife speed was fixed at 900 rpm with an 8.38 mm die opening. Kibbles were dried in a convection oven (Model FP 240; Binder Inc.; NY, U.S.A.) at 100 ºC until moisture content was below 10%. There was no coating step because these treatments as diets were not intended to be fed to dogs.

During extrusion, a total of 20 samples were collected at the exit of the extruder barrel (off the extruder; OE) once production reached steady state (after 5 minutes of changing the settings), immediately flash frozen in liquid nitrogen, and kept at -70°C until RVA analysis was performed. Samples were placed into 30 g plastic bags (Whirl-Pak, Madison, WI) and collected at two time points during each run to assure representative sampling, which were then used for RVA and starch cook analyses. Samples collected off the drier (OD) were used for kibble measurements, texture analysis, and resistant starch determination.

Output parameters collected at the extruder panel were motor load, PC temperature, pressure and extrudate temperature measured by a penetrating probe between the die insert and final head of the barrel, while in-barrel moisture (IBM) and specific mechanical energy (SME) were considered as intermediate variables (Table 3). The SME was determined according to Equation 1:

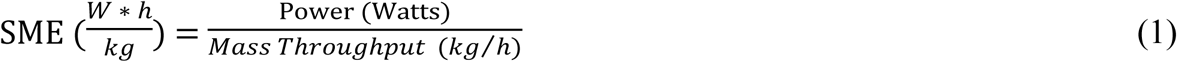

Where motor power was measured by the solid-state extruder motor drive, and mass throughput was calculated as the total wet mass flow rate (dry feed rate + water and steam inputs at PC and extruder). The steam input was corrected for calculated steam loss before the extruder. In-barrel moisture was determined according to Equation 2:

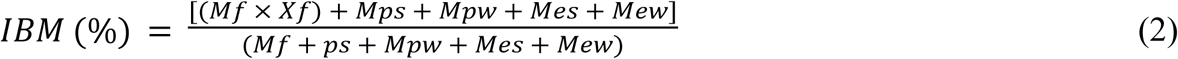

Where: Mf = dry feed rate (kg/h); Xf = wet basis moisture content of the feed material (%); Mps = rate of steam condensation onto raw material blend (kg/hr); Mpw= water injection rate in the pre-conditioner (kg/hr); Mes = steam injection rate in the extruder (kg/hr), and Mew= water injection rate in the extruder (kg/hr).

### 2.5 Kibble measurements and texture analysis

Kibble dimension measurements were performed on 20 kibbles per sample. Each kibble was randomly selected, then diameter and length were each measured twice with a digital caliper, averaged, and weighed on an analytical balance (Ohaus, Explorer: E1RW60, OHAUS, Parsippany, NJ). With this information, piece density (g/cm^3^) and sectional expansion index (SEI; cm^2^_e_ /cm^2^_d_) were calculated. Volumetric expansion index (VEI) was calculated according to (12):

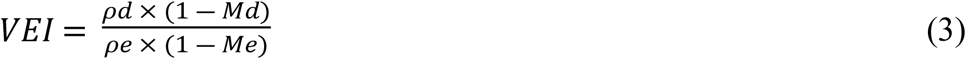

Where: ρd = extrudate density inside the die; Md= moisture content of the extrudate in the die; ρe= apparent density of the wet kibble; and Me= moisture content of the wet kibble. Moisture content inside the die (Md) equaled IBM, while moisture content of the extrudate after exiting the die (Me) was calculated as IBM minus steam loss. Steam loss was estimated according to (13). Density of the kibble inside the die (ρd) was calculated using a model (14). Lastly, longitudinal expansion index (LEI) was calculated as a function of VEI divided by SEI.

Kibble hardness was considered the peak force required to break the kibble at the primary significant fracture. This was determined on 30 kibbles per run using a texture analyzer (TA-XT2; Texture Technology Corp., Scarsdale, NJ, U.S.A.) equipped with a 50 kg load cell. A 25 mm cylindrical probe was used to compress kibbles at a 50% strain level.

### 2.6 Starch analyses

Kibbles were subjected to RVA analysis as an indicator of process effects on starch biopolymers. Briefly, uncoated wet kibbles (OE) were frozen at -70°C until analysis, and then ground through a meat mincer and sieved using a 400 mm sieve screen size. Prior to the RVA analysis, sample moisture was determined, then 2 g was diluted with 25 g deionized distilled water in an aluminum cup containing a plastic paddle. The RVA (RVA 4800; Perten and PerkinElmer Instruments, Springfield, IL) was performed for 23 min. Samples were equilibrated at 25°C for 2 min at 960 rpm, then the speed decreased to 160 rpm and temperature increased to 95°C between 2 and 10 min, then held at 95°C for 3 minutes then decreased to 25°C from 13 to 18 min. The RVA data are reported as area under the curve (AUC) for each peak (cold peak, raw peak and setback). The ratio of raw:cooked starch was also calculated by dividing the AUC of the raw peak by the AUC of the cold peak.

Resistant starch determinations were performed according to an enzymatic assay with glucose measured by colorimetry (K-RSTAR; Megazyme International Ireland Limited, Ireland). Degree of gelatinized starch was determined by a modified glucoamylase test based on a 70-minute enzymatic hydrolysis (15) with a blend of amylase and amyloglucosidase from *Aspergillus niger* (Hazyme DCL; DSM corporation, Heerlen, Netherlands), and glucose quantified by a YSI Model 2700 glucose analyzer (YSI; Yellow Springs, OH, U.S.A.).

### 2.7 Statistical analysis

The surface response model of central composite data from small scale (Evolum 25) extrusion was first analyzed for lack of fit with the RSREG procedure from Statistical Analysis Software (SAS; v. 9.4, SAS Institute Inc., Cary, NC, USA). When lack of fit was not significant (*P* > 0.05) data were analyzed by the REG procedure (SAS; v. 9.4, SAS Institute Inc., Cary, NC, USA). The main effects of PS, extruder shaft speed (SS), and IBM were added to the model, as well as all cross-products and quadratic terms. The model with the best fit (lowest P-value) was obtained by backwards elimination. When a main effect was eliminated from the model, but either cross-product or quadratic term were significant, the main effect was added back. The intercept was kept in the model even when it was not significant. This procedure was applied to RS, starch cook and RVA as dependent variables. When a model was significant (*P* < 0.05) with an adjusted R^2^ > 0.50, a surface response plot was created using the proc G3D procedure from SAS (SAS 9.4, Cary NC). Pearson correlations were performed between starch measurements, kibble data and extrusion parameters using the CORR procedure from SAS (SAS v 9.4, Cary, NC). Additionally, two regression equations were constructed using the REG procedure from SAS between RS and SME, and RS and dough T at the end of the extruder barrel.

## 3. Results

### 3.1 Particle size

The median circle equivalent (CE) diameter distribution measured by the Morphologi G3 for the three ground corn samples were between 3.6 and 5.5 times lower than the sieve sizes used to grind each corn (Table 2), and high sensitivity (HS) circularity, aspect ratio and elongation were numerically similar within ground corn and dry mixes. Particle size distributions were bimodal for corn ground at the three levels (Fig 1). Corn ground using a 0.793, 1.19, and 1.586 mm sieve size had a mean geometric diameter ± standard deviation of 158.6 ± 2.05, 174.8 ± 2.24 and 221 ± 2.6 μm, respectively, and the corresponding particle sizes of the dry mix (corn mixed and ground with other ingredients) were 161.6 ± 1.99, 172.7 ± 2.08 and 195.3 ± 2.24 μm.

**Table 2.** Particle size analyses determined by the Morphologi G3 Instrument and the Ro-Tap Procedure.

| Analysis | Corn 1 | Corn 2 | Corn 3 | Dry mix 1 | Dry mix 2 | Dry mix 3 |
| --- | --- | --- | --- | --- | --- | --- |
| Morphologi G3 Instrument |  |  |  |  |  |  |
| Number of particles analyzed | 76,892 | 51,448 | 41,324 | 24,479 | 30,924 | 29,871 |
| <sup>1</sup>CE diameter distribution (v, 0.5), $\mu\text{m}$ | 183.4 | 331.6 | 287.3 | 152.9 | 151.1 | 160.4 |
| <sup>2</sup>HS circularity (n, 0.5) | 0.879 | 0.901 | 0.913 | 0.781 | 0.798 | 0.808 |
| <sup>3</sup>Aspect ratio number distribution (n, 0.5) | 0.782 | 0.796 | 0.806 | 0.740 | 0.742 | 0.745 |
| <sup>4</sup>Elongation (n, 0.5) | 0.215 | 0.201 | 0.191 | 0.258 | 0.256 | 0.253 |
| Ro-tap procedure |  |  |  |  |  |  |
| Mean geometric diameter (Dgw), $\mu\text{m}$ | 158.6 | 174.8 | 221.1 | 161.6 | 172.7 | 195.3 |
| Standard deviation (Sgw) | 2.05 | 2.24 | 2.6 | 1.99 | 2.08 | 2.24 |

Corn 1, 2 and 3 ground with a 0.793, 1.19 and 1.59 mm screen size, respectively.
Dry mix 1, 2 and 3 were the same diet recipe mixed with corn 1, 2 and 3, respectively.
<sup>1</sup>CE= circle equivalent; (v, 0.5) = median of the volume distribution. The volume is calculated by converting average 3D image of the particles to a 2D circle of equivalent area.
<sup>2</sup>HS= high sensitivity circularity; (n, 0.5) = median of the number distribution. Calculated as $4\pi A/P^2$ ; where A is the particle area, and P its perimeter. Measurements ranged from 0 to 1 and described the deviation from a perfect circle (the closer to 1, the more it approximates to a perfect circle).
<sup>3</sup>Aspect-Ratio-Number Distribution; (n, 0.5) = median of the average width/length of the particles. It ranges from 0 to 1. A low aspect-ratio would assume a rod shape.
<sup>4</sup>Elongation= average particle elongation (the closer to 1, the longer it is). Calculated as $1 - \text{aspect-ratio}$ (ranges from 0 to 1).

**Fig 1.**
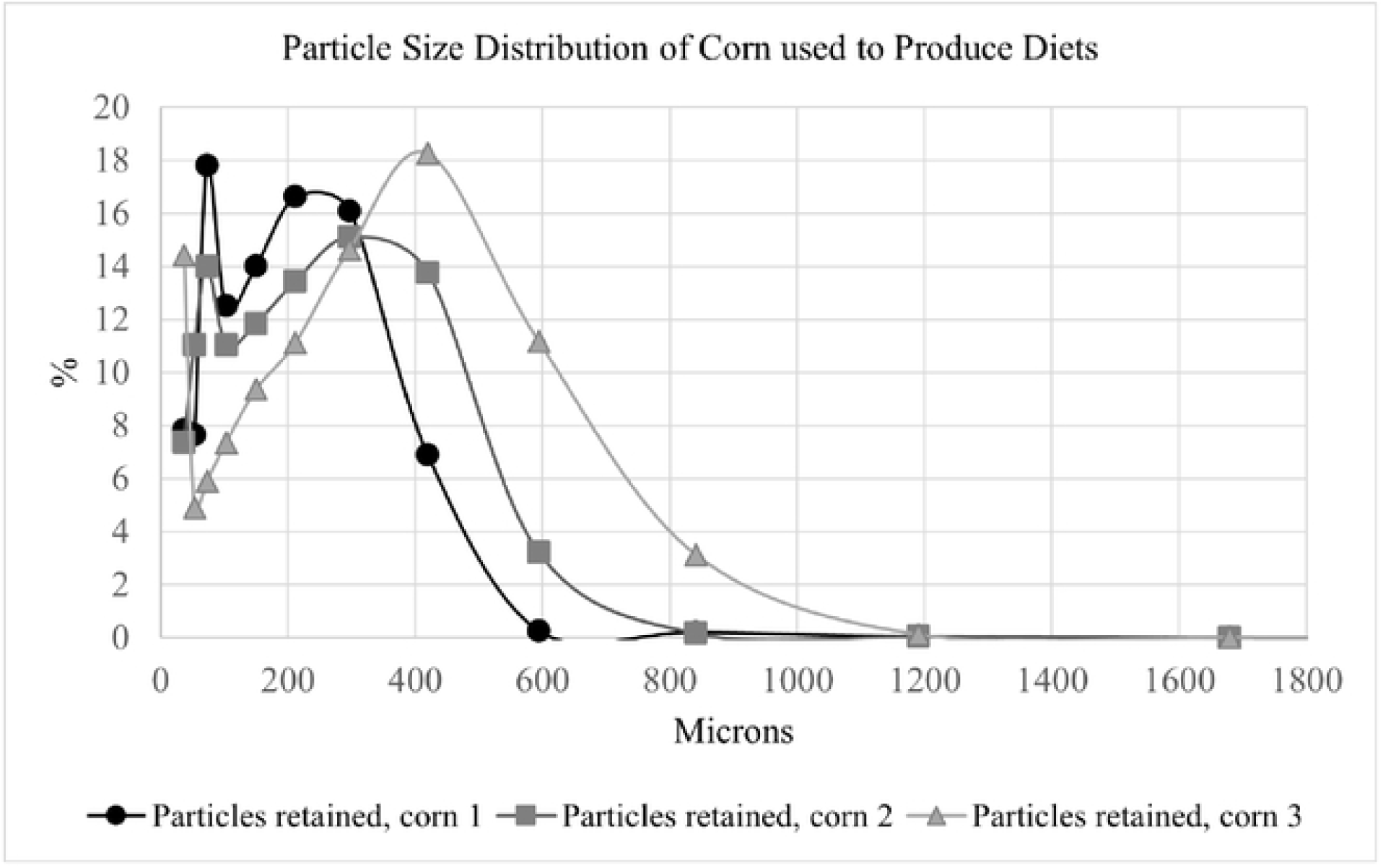
Particle Size Distribution of Corn Ground in a Hammermill with Sieve Sizes 0.79, 1.19 and 1.59 mm (Corn 1, 2 and 3, respectively).

### 3.2 Extrusion processing

The intermediate extrusion parameters IBM and SME were targeted to have the greatest variation according to each treatment combination. The 6 replicates had an IBM of 32.4%, while SME varied from 30.2 to 32.5 Wh/kg (Table 3). In-barrel moisture was the lowest on treatments 12, 11, 19, 16 and 17 (average 29.8% ± 0.87), and the highest in treatments 14, 15, 9, 19 and 20 (average 35.3% ± 0.70). Specific mechanical energy was intentionally targeted to change according to water additions and shaft speed modifications, ranging from 11.6 to 52 Wh/kg (Table 3). Treatments with the highest SME (> 50 Wh/kg) were those with the high extruder shaft speed setting, and either low or intermediate IBM. Conversely, the lowest SME values (11.6, 15.7 and 12.8 Wh/kg) were obtained with the low extruder shaft speed setting, and either high or intermediate IBM.

**Table 3.** Input, Intermediate and Output Parameters recorded at the Evolum 25 Extrusion Panel.

| Sample | Input parameters<sup>1</sup> |  |  | Intermediate<sup>2</sup> and output<sup>3</sup> parameters |  |  |  |  |  |
| --- | --- | --- | --- | --- | --- | --- | --- | --- | --- |
|  | PC water,<br>kg/h | PC steam,<br>kg/h | Feed<br>Moisture, % | Motor<br>Load, % | PC Temp.,<br>°C | IBM, % | SME,<br>Wh/kg | Extrudate<br>Temp. at the<br>Die, °C | Extrudate<br>Pressure at<br>the Die, bar |
| 1 | 3 | 4.5 | 11.05 | 8 | 80 | 32.4 | 32.5 | 118 | 11.0 |
| 2 | 3 | 4.5 | 11.05 | 8 | 81 | 32.4 | 31.6 | 116 | 11.0 |
| 3 | 3 | 4.5 | 11.05 | 8 | 80 | 32.4 | 31.2 | 117 | 11.0 |
| 4 | 3 | 4.5 | 11.05 | 8 | 83 | 32.4 | 30.5 | 118 | 11.0 |
| 5 | 3 | 4.5 | 11.05 | 8 | 85 | 32.4 | 30.3 | 116 | 11.0 |
| 6 | 3 | 4.5 | 11.05 | 8 | 83 | 32.4 | 30.2 | 117 | 11.0 |
| 13 | 3 | 4.3 | 11.05 | 8 | 87 | 32.1 | 30.0 | 116 | 11.0 |
| 12 | 1.5 | 4.3 | 10.93 | 8 | 88 | 29.3 | 51.8 | 128 | 10.5 |
| 11 | 1.5 | 4.1 | 10.93 | 11 | 88 | 28.9 | 22.6 | 118 | 17.0 |
| 14 | 4.5 | 4.5 | 10.93 | 6 | 86.5 | 34.8 | 34.9 | 116 | 8.0 |
| 15 | 4.5 | 4.4 | 10.93 | 8 | 88 | 34.7 | 11.6 | 108 | 11.0 |
| 7 | 3 | 4.45 | 10.93 | 8.5 | 85 | 32.3 | 51.2 | 127 | 10.5 |
| 8 | 3 | 4.5 | 11.05 | 9.5 | 87 | 32.4 | 15.7 | 113 | 13.5 |
| 9 | 4.5 | 4.5 | 11.05 | 6 | 86 | 34.9 | 22.3 | 114 | 9.0 |
| 10 | 1.5 | 4.2 | 11.05 | 8.5 | 88.5 | 29.2 | 34.1 | 117 | 12.0 |
| 16 | 1.5 | 4.9 | 11.13 | 8 | 85.5 | 30.6 | 52.0 | 130 | 10.5 |
| 17 | 1.5 | 5 | 11.13 | 8 | 86 | 30.8 | 20.3 | 118 | 17.0 |
| 18 | 3 | 4.9 | 11.13 | 8 | 86 | 33.2 | 32.3 | 119 | 12.0 |
| 19 | 4.5 | 5.3 | 11.13 | 8 | 86 | 36.2 | 35.0 | 118 | 8.0 |
| 20 | 4.5 | 5.1 | 11.13 | 8 | 85.5 | 35.9 | 12.8 | 108 | 11.5 |

<sup>1</sup>The following parameters were kept constant across all treatments: PC shaft speed set at 100 kg/h; feed rate at 30 kg/h; extruder water and steam at 2 and 0 kg/h, respectively, and knife speed at 900 rpm.
<sup>2</sup>Intermediate parameters: IBM and SME.
<sup>3</sup>Output parameters: motor load, PC temperature, extrudate temperature at the die and Extrudate pressure at the die.

### 3.3 Kibble measurements and starch analyses

Kibble volume and density for the 20 treatments ranged from 0.995 to 1.429 cm^3^ (average 1.21 cm^3^ ± 0.125) and 0.377 to 0.732 g/cm^3^ (average 0.463 g/cm^3^ ± 0.076), respectively (S. Table 2). Kibble expansion (SEI) ranged from 1.328 to 2.082 times the die size (average 1.738 ± 0.2109), and VEI and LEI ranged from 0.719 to 1.68 cm^3^_e_ /cm^3^_d_ (average 1.27 ± 0.228), and 0.385 to 0.961 cm^2^_e_ /cm^2^_d_ (average 0.732 ± 0.119), respectively. Volumetric expansion index (VEI) represents the overall kibble expansion, while SEI and LEI are two components of expansion. Finally, hardness ranged from 6.37 to 16.41 kg (average 10.15 kg ± 2.861; S. Table 2).

Although the 20 samples were produced with the same basal recipe, their total starch content varied from 48.8 to 65.9% (mean 57.7% ± 4.25; Table 4). This was likely due to the high variability intrinsic to the total starch assay. Starch cook of the raw dry mixes with fine, medium and coarsely ground corn were, respectively, 11.0, 10.7 and 9.5%, and after processing these ranged from 83.3 to 99.7%; (mean 91.5% ± 5.13; Table 4). The RS content of the raw dry mixes with fine, medium and coarsely ground corn were 1.20%, 1.05% and 2.28%, respectively, and RS of the diets after extrusion processing varied from 0.24 to 1.48% (mean 0.79 ± 0.278; Table 4).

**Table 4.** Starch Analyses of Samples produced through the Small-Scale Twin-Screw Extruder (Evolum 25).

| Sample | Total Starch<sup>1</sup>, % | Starch Cook, % | RVA Cooked Starch AUC<sup>2</sup> | RVA Raw Starch AUC<sup>2</sup> | RVA Setback AUC<sup>2</sup> | RVA Raw:cooked<sup>4</sup> | Resistant Starch<sup>3</sup>, % |
| --- | --- | --- | --- | --- | --- | --- | --- |
| 1 | 57.9 | 83.3 | 967 | 2,340 | 8,182 | 2.42 | 0.821 |
| 2 | 61.2 | 89.4 | 1,078 | 2,084 | 7,302 | 1.93 | 0.731 |
| 3 | 59.3 | 92.9 | 1,055 | 1,816 | 5,963 | 1.72 | 0.737 |
| 4 | 61.6 | 87.0 | 928 | 2,044 | 6,508 | 2.20 | 0.820 |
| 5 | 59.5 | 88.3 | 2,451 | 1,864 | 6,984 | 0.76 | 0.928 |
| 6 | 55.3 | 92.9 | 1,035 | 2,307 | 6,953 | 2.23 | 0.860 |
| 7 | 54.6 | 93.2 | 1,089 | 2,279 | 7,422 | 2.09 | 0.240 |
| 8 | 51.1 | 90.7 | 1,186 | 2,137 | 8,345 | 1.80 | 1.057 |
| 9 | 55.0 | 83.6 | 1,183 | 1,992 | 9,492 | 1.68 | 0.952 |
| 10 | 48.8 | 99.2 | 1,189 | 2,051 | 5,575 | 1.72 | 0.751 |
| 11 | 52.5 | 93.6 | 1,228 | 2,443 | 8,170 | 1.99 | 0.721 |
| 12 | 61.7 | 99.7 | 1,748 | 1,336 | 2,590 | 0.76 | 0.326 |
| 13 | 65.9 | 90.1 | 2,479 | 2,233 | 6,060 | 0.90 | 0.573 |
| 14 | 60.3 | 88.6 | 1,087 | 2,497 | 7,614 | 2.30 | 0.719 |
| 15 | 52.7 | 94.0 | 759 | 2,131 | 10,558 | 2.80 | 0.903 |
| 16 | 59.9 | 99.6 | 3,239 | 1,199 | 2,915 | 0.37 | 0.350 |
| 17 | 59.0 | 84.6 | 1,037 | 1,995 | 7,962 | 1.92 | 1.056 |
| 18 | 57.2 | 98.8 | 1,243 | 1,789 | 6,227 | 1.44 | 0.907 |
| 19 | 61.1 | 92.9 | 1,364 | 1,740 | 4,560 | 1.28 | 0.801 |
| 20 | 59.3 | 88.0 | 850 | 2,487 | 9,392 | 2.93 | 1.480 |

Viscosity (RVA) measurements were determined on wet kibbles (out of the extruder), and resistant starch, total starch and starch cook were determined on dried kibbles.
<sup>1</sup>Starch cook of Dry mix 1, 2 and 3 were 11.0, 10.7 and 9.5%, respectively.
<sup>2</sup>Expressed as relative viscosity units (RVU)
<sup>3</sup>Resistant starch of Dry mix 1, 2 and 3 were 1.20, 1.05 and 2.28%, respectively.

Each stage of change in viscosity measured by RVA was calculated as follows: The area under the curve (AUC) between minute 0.4 and 6.0 was the cooked starch AUC; the AUC between 6.1 and 14 minutes represented the raw starch AUC, while setback viscosity (high molecular weight starch) was measured as the AUC between minutes 14.1 and 23. Although starch is responsible for the majority of viscosity changes, other nutrients such as protein or fiber may interfere with the analysis, thus in the present study peak viscosities were not relevant and not reported. To illustrate the RVA curves, two samples with extreme extruder barrel temperatures at the end of the barrel were plotted (Fig 2). The lower energy sample (number 15) showed little to no initial viscosity, a pronounced native starch peak where the RVA temperature increased above 60ºC, and a high setback viscosity. In contrast, the sample produced at higher energy (number 16) exhibited a large initial peak and no indication of a native starch (raw starch) peak where the RVA temperature increased. The setback viscosity of sample 16 was lower than sample 15. The ratios between the RVA raw:cooked AUC were calculated as another means to estimate the extent of starch cooked relative to raw starch, and these were 2.80 and 0.37 in extreme samples 15 and 16, respectively (mean 1.76 ± 0.681; Table 4).

**Fig 2.**
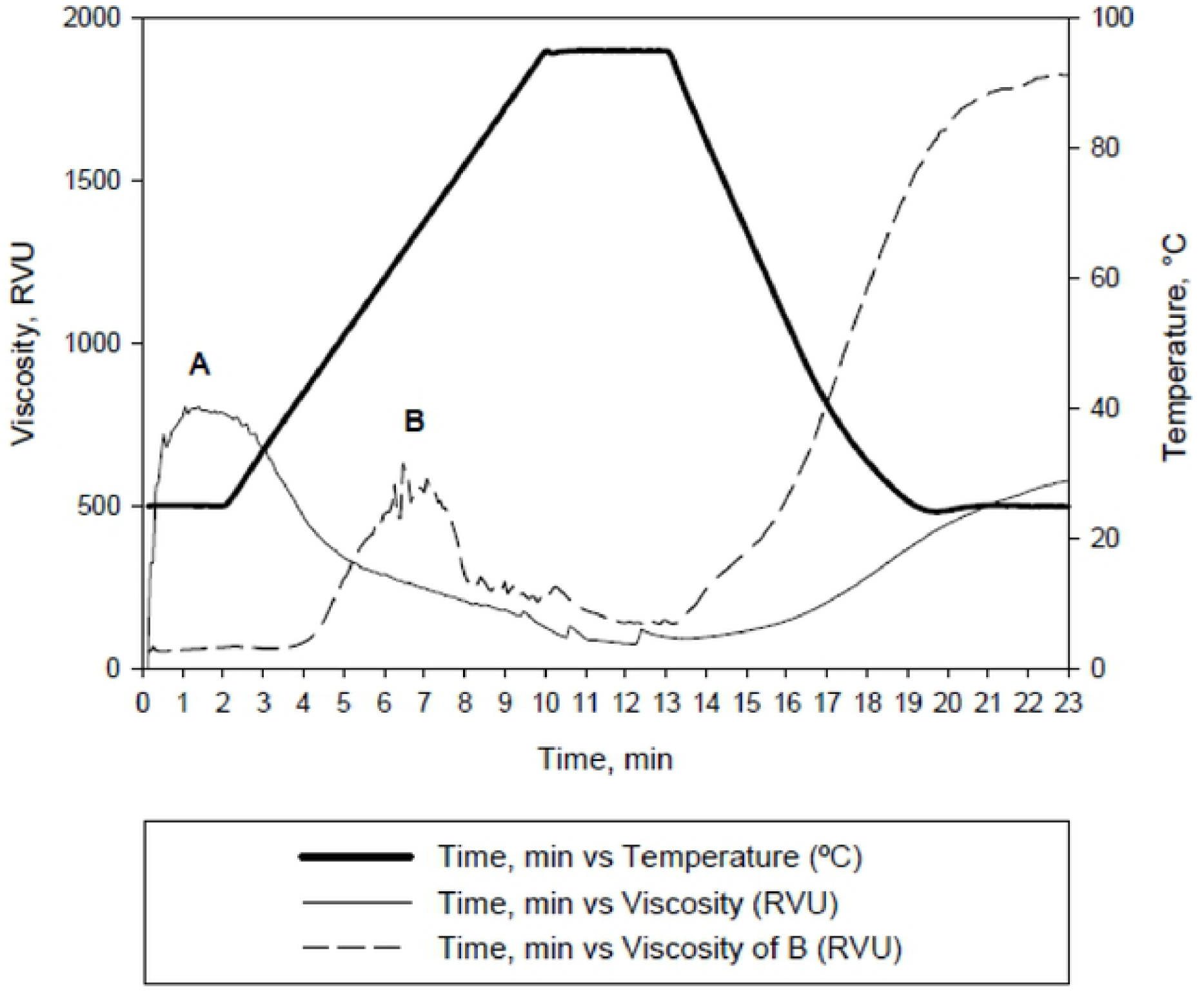
Viscosity Curves measured by RVA of Wet Kibbles of a High (Sample 16, A) and a Low (Sample 15, B) Thermomechanical Energy Process.

### 3.4 Correlations between starch analyses and processing parameters

Resistant starch was less than 50% negatively correlated (*P* < 0.05) with starch cook and RVA cooked AUC, and also less than 50% positively correlated (*P* < 0.05) with RVA raw:cooked AUC and setback viscosity (Table 5). Conversely, the starch percent cook results had an inverse relationship with RVA raw AUC, raw:cooked AUC, and setback viscosity. Setback viscosity had the greatest correlations with RS and starch cook among other RVA parameters, indicating that the greater the content of native starch, the greater the final viscosity due to the presence of longer amylose and amylopectin molecules.

**Table 5.** Pearson Correlation between Chemical and Physical Methods of Starch Analyses.

| Item |  | Resistant Starch | % Starch Cook |
| --- | --- | --- | --- |
| % Cook | *r* | -0.498 | - |
|  | *P* | 0.026 | - |
| RVA cooked starch AUC | *r* | -0.452 | 0.324 |
|  | *P* | 0.046 | 0.164 |
| RVA raw starch AUC | *r* | 0.418 | -0.525 |
|  | *P* | 0.066 | 0.017 |
| RVA Raw: cooked ratio | *r* | 0.519 | -0.420 |
|  | *P* | 0.019 | 0.065 |
| RVA Setback viscosity AUC | *r* | 0.632 | -0.624 |
|  | *P* | 0.003 | 0.003 |

Volumetric expansion index (VEI) had a strong negative correlation with IBM (r=-0.89), which meant that a lower moisture content led to greater overall kibble expansion within the experimental parameters (Table 6). Longitudinal expansion index (LEI) was mostly affected by dough temperature (r=0.61) and IBM (r=-0.733), while SEI did not have a significant correlation with any of the parameters tested.

**Table 6.** Pearson Correlation between Kibble Endpoints and Extrusion Parameters.

| Item |  | VEI, | LEI | SEI | RS | %<br>Starch<br>Cook | RVA,<br>Cooked<br>AUC | RVA,<br>Raw AUC | Raw:<br>Cooked<br>ratio | Setback<br>viscosity<br>AUC |
| --- | --- | --- | --- | --- | --- | --- | --- | --- | --- | --- |
| Dough T at end | r | 0.396 | 0.610 | -0.179 | -0.829 | 0.475 | 0.507 | -0.603 | -0.615 | -0.772 |
| of barrel, °C | P | 0.084 | 0.004 | 0.450 | <.0001 | 0.035 | 0.023 | 0.005 | 0.004 | <.0001 |
| Shaft speed, rpm | r | -0.076 | 0.162 | -0.311 | -0.724 | 0.327 | 0.394 | -0.448 | -0.494 | -0.694 |
|  | P | 0.749 | 0.495 | 0.182 | 0.000 | 0.160 | 0.086 | 0.048 | 0.027 | 0.001 |
| IBM, % | r | -0.890 | -0.733 | -0.350 | 0.482 | -0.408 | -0.305 | 0.266 | 0.384 | 0.433 |
|  | P | <.0001 | 0.000 | 0.130 | 0.031 | 0.075 | 0.191 | 0.257 | 0.094 | 0.056 |
| SME, Wh/kg | r | 0.302 | 0.476 | -0.159 | -0.862 | 0.486 | 0.485 | -0.538 | -0.580 | -0.800 |
|  | P | 0.196 | 0.034 | 0.504 | <.0001 | 0.030 | 0.030 | 0.014 | 0.007 | <.0001 |
| PS, Dgw | r | -0.232 | -0.046 | -0.241 | 0.342 | 0.039 | 0.101 | -0.327 | -0.117 | -0.177 |
|  | P | 0.324 | 0.847 | 0.306 | 0.140 | 0.870 | 0.671 | 0.159 | 0.623 | 0.456 |

VEI= volumetric expansion index; LEI= longitudinal expansion index; SEI= sectional expansion index; RS= resistant starch; IBM= in-barrel moisture; SME= specific mechanical energy; PS= particle size.

Resistant starch had a strong negative correlation with dough temperature at the end of the barrel, shaft speed and SME (r= -0.829, -0.724 and -0.862, respectively), and a less strong positive correlation with IBM (r= 0.48; Table 6). The regression equations of interest follow below:

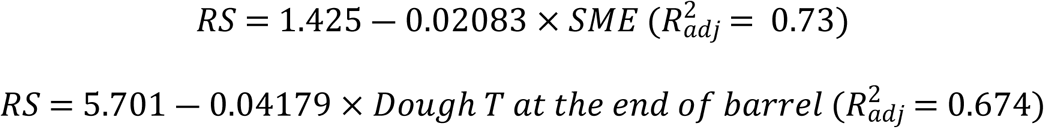

Starch cook had a low correlation with all parameters tested but was significant (*P* < 0.05) and positively correlated with dough temperature and SME and negatively correlated with IBM. The RVA AUC of cooked and raw starches were mostly affected by dough temperature and SME, but not IBM. Lastly, setback viscosity had a strong negative correlation with dough temperature and SME.

### 3.5 Surface response plots and model building

Dry kibble RS was the primary endpoint. For model building, the actual measured rather than target parameters were used. The independent variables were PS at 161.5 um, 172.7 um and 195.3 um mean geometric diameter, IBM at 29.8, 32.5 and 35.3%, and SS at 400, 800 and 1200 rpm, representing the low, medium and high settings. The IBM obtained in each level had to be averaged in order to be used in the model building. The model using these variables as predictors for RS content in the dry kibble was linear (*P* < .0001) with non-significant lack of fit (*P* = 0.1136). After backward elimination and regression analysis testing the main effects, their cross-products and quadratic terms the resultant significant terms were SS (*P* = 0.1439), PS (*P* = 0.0033), IBM (*P* = 0.0003), and SS*PS (*P* = 0.0427). The final model (*P* < .0001) with an adjusted R^2^ of 0.834, MSE 0.0129, 11.04% coefficient of variation (CV), was:

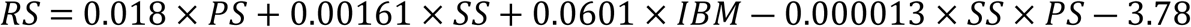

Due to difficulties visualizing a 3D surface response plot using 3 independent variables (PS, SS, and IBM), two variables were plotted against RS for each (Figs 3-5).

**Fig 3.**
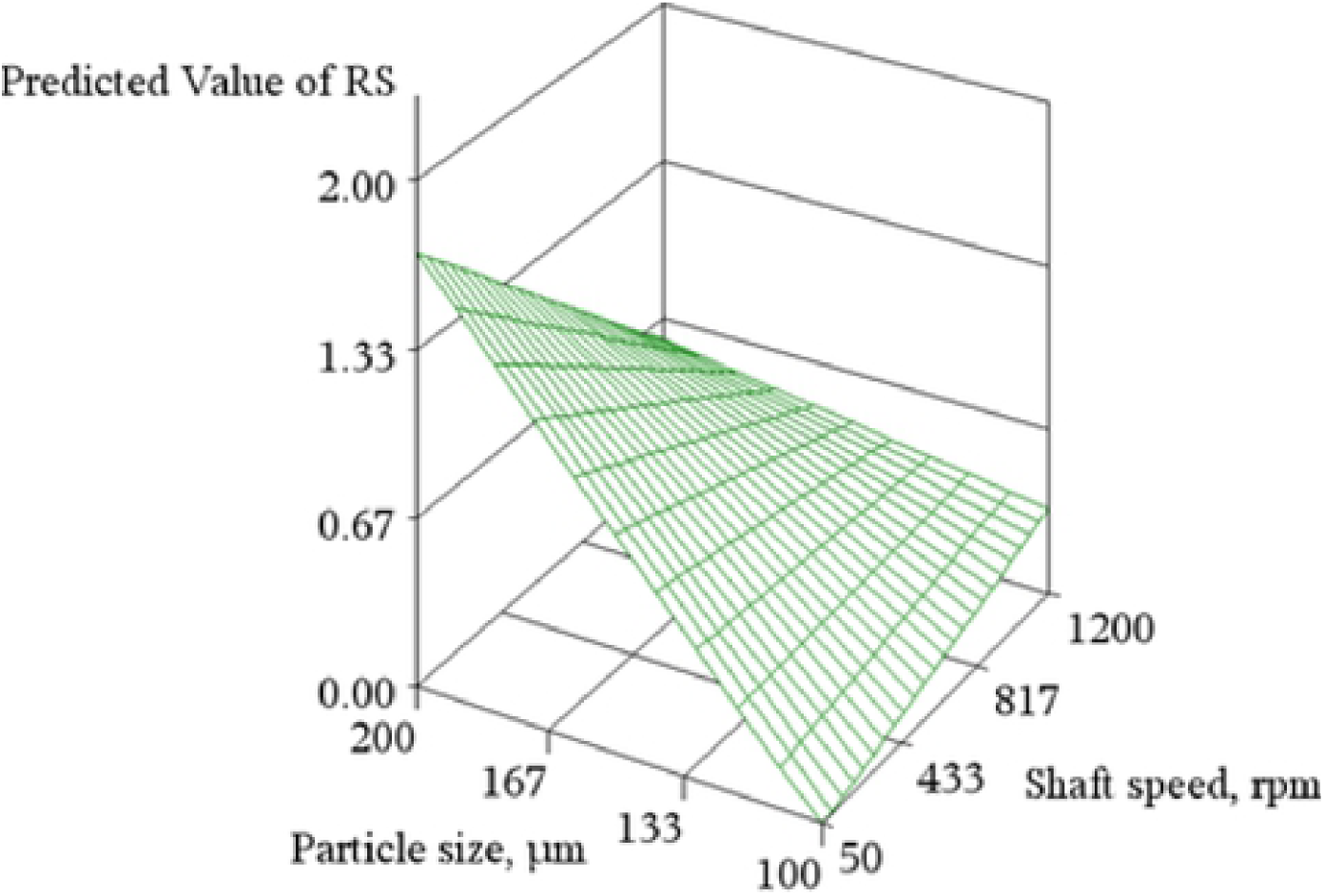
Surface Response Plot using Particle Size and Extruder Shaft Speed to predict Resistant Starch Levels in an Extruded Dog Food.

**Fig 4.**
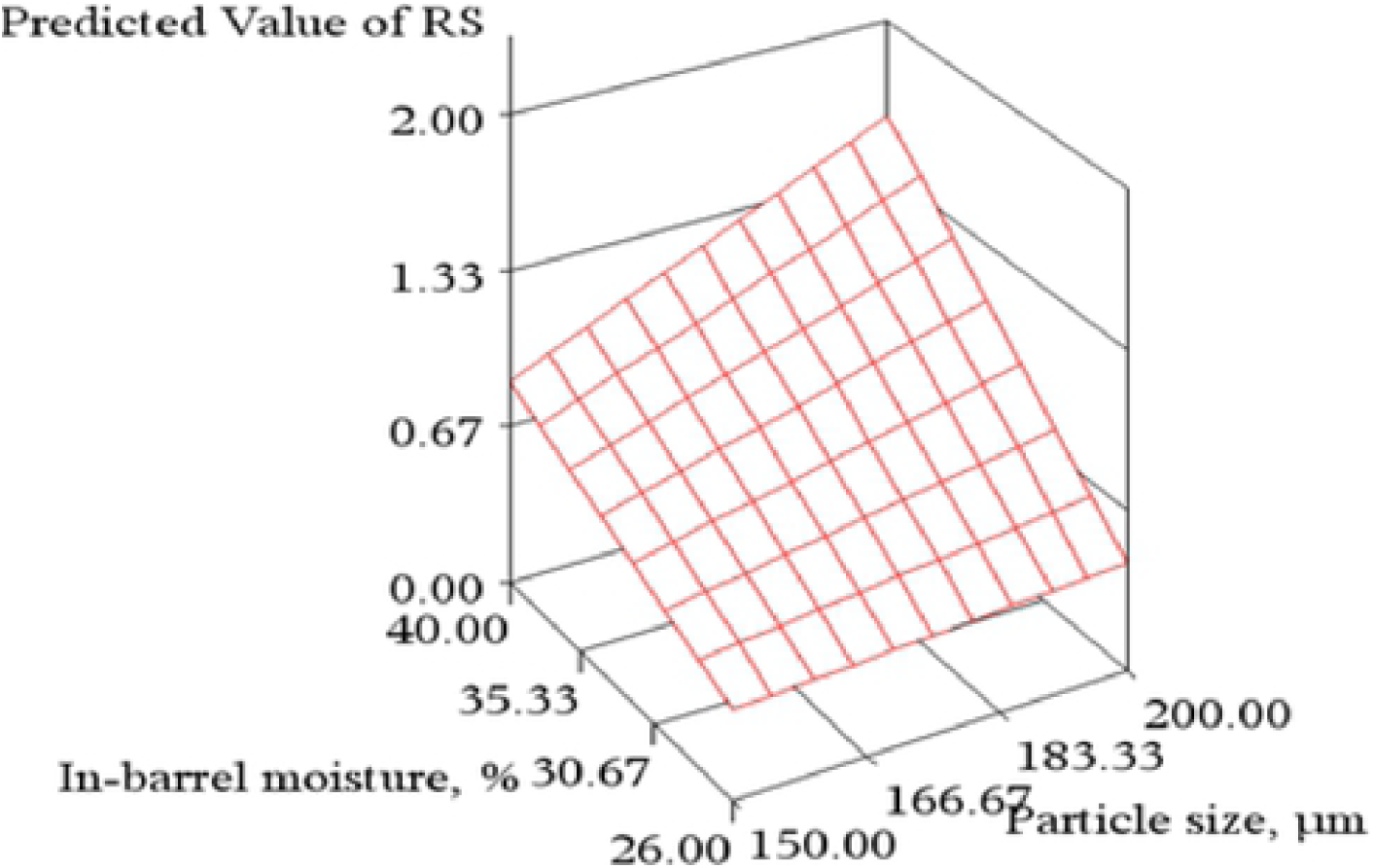
Surface Response Plot using In-Barrel Moisture and Particle Size to predict Resistant Starch Levels in an Extruded Dog Food.

**Fig 5.**
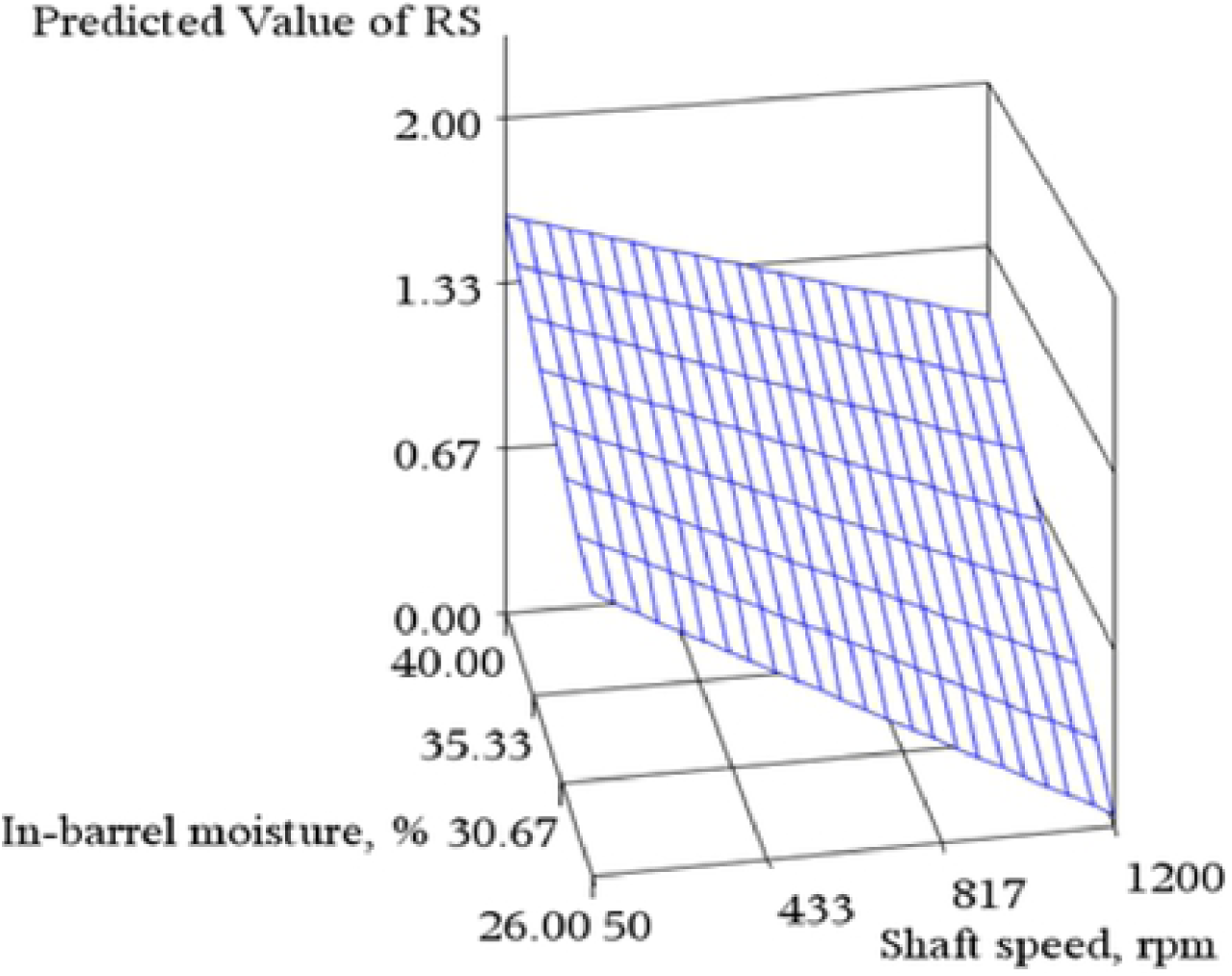
Surface Response Plot using In-Barrel Moisture and Extruder Shaft Speed to predict Resistant Starch Levels in an Extruded Dog Food.

The final model to predict viscosity (RVA) with endpoints cooked peak and raw:cooked ratio was not significant (*P* > 0.05). The model to predict RVA raw peak AUC was significant (*P* = 0.0084) after backwards elimination with IBM, SS, PS and IBM*SS as part of the final model; but the graphs were not created because the adjusted R^2^ was low (0.464) with CV=12.5% and mean AUC 24,458:

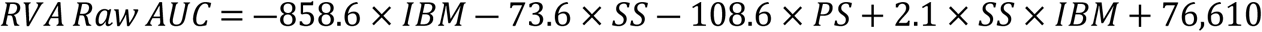

Gelatinized starch (% cook) had a non-significant lack of fit (*P* = 0.2509), and the model was significant (*P* = 0.0401) after backwards elimination. However, the predictor IBM was not significant (*P* = 0.4779) and both SS and their cross-product (SS*IBM) had only a tendency (0.10 > *P* > 0.05) to be significant. The final model also had a low R^2^adj (0.283):

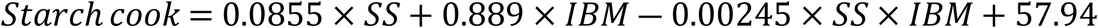

## 4. Discussion

To date, the present study is the first to determine the effect of pet food processing parameters on RS yield and to create a model that would predict RS content in a dry extruded kibble. A secondary goal was to explore methods of starch gelatinization, correlate these among themselves, as well as correlate extrusion processing parameters with starch transformation measures and kibble parameters.

The RS targeted in the present study was likely a combination of types II and III (16,17). Recently, a broader classification was suggested which identifies this type of RS as starches not digestible by enzymes due to their tightly packed crystalline structure (18) which is present in raw starches (19).

The ideal concentration of RS that benefits dog colonic health without decreasing stool quality still hasn’t been established for all breeds. Although it is known that large dogs may produce loose stools when fed a diet with RS as low as 2.5% (8). In the present study the raw dry mix with the most RS (dry mix 3) had only 2.2% RS. Thus, the maximum RS in a treatment could not have been over 2.2%. Ideally, we would extrude a dog food with the minimal energy necessary to destroy antinutritional factors and pathogens, while minimizing starch cook. This is challenging since extruded pet foods need hydration as well as thermal and mechanical energies to form an expanded kibble.

Starch granules of cereals like corn, wheat or rice have an X-ray diffraction type A and also contain pores and channels that facilitate alpha-amylase adhesion and digestion of the substrate (18,20,21). Conversely, tubers have large smooth starch granules with less enzyme adhesion sites, and legume starches are trapped within cotyledon cells, which are little disrupted after thermal processing (22). Although tubers and legumes tend to be higher in naturally occurring RS than cereals (23), corn was chosen as the sole starch ingredient for two reasons: 1.) it is one of the most common cereal grains used in pet food due to its high apparent total tract digestibility coefficients, palatability and low cost. 2.) by choosing one starch source and not adding any fiber ingredients the effect of recipe would be minimized and the focus would be on the effects of processing on RS yield. Moreover, whole ground corn was selected instead of corn flour because it has lower *in vitro* starch digestibility when compared to fine milled flours (24).

It is well known that raw cereals, legumes or tubers possess a greater amount of RS than their cooked forms (25,26), but thermal processing is usually required to destroy antinutritional factors (27), increase acceptability (25), and microbiological safety of products. Thermal and mechanical processing with some moisture gelatinizes the starch making the α-glucan chains less ordered and more available for enzymatic adsorption and digestion (28,29). The RS of kibbles in the present study had a high and inverse correlation with dough temperature at the end of the extruder barrel confirming that the lowest thermomechanical energy from the process resulted in the most RS. There are many processing inputs that contribute to changes in extrudate temperature such as feed rate, steam and water additions, extruder shaft speed, and die open area.

When corn is extensively ground, starch gelatinization and digestion are improved due to a large surface area to mass of the small particles. This happens because the starch ground to smaller particles becomes more exposed to hydration, which increases gelatinization. Conversely, coarsely ground corn has a lower surface area to mass which slows water penetration and starch gelatinization, thereby decreasing its digestion (4,5). Grain milling also destroys cell walls and disrupts the protein matrix making starch more available (30). For these reasons, an important factor selected to construct the surface response model for RS in dry extruded kibbles was the corn grind size. In the present study, the largest mean geometric diameter was less than initially targeted. This likely happened because the hammermill used was more effective than expected, and the corn (and the ration it was contained within) was ground through the hammermill with a 1.59 mm screen size twice. A better model may have been created if wider differences among mean geometric diameter had resulted. In future work using a larger screen size would be recommended. (5) and (4) reported that extruded dog food produced with raw corn ground to geometric mean diameter of 224 (low) and 312 μm (high) and at high and low SME inputs yielded 0.21-0.22 and 1.46-1.54% RS, respectively. In their work the lowest geometric mean was greater than the grain-mix coarsely ground in the present study. Moreover, in their studies the SME was controlled by differing the die open area rather than altering screw speed like in the present work, and both methods were effective in controlling SME. (31) reported that fine, medium and coarse maize (360, 452 and 619 μm mean geometric diameter, respectively) produced foods with a starch gelatinization of 79.9, 73.8 and 63.2%, respectively. The starch cook values they obtained were wider than ours due to the greater differences in particle size. While RS was not measured in that work (31), it would likely be above what was determined in the present study. Nevertheless, particle size still had a significant effect on RS concentration when plotted against shaft speed and in-barrel moisture; wherein, the RS prediction was the greatest at the largest mean geometric diameter.

Water, which is required for starch gelatinization, has a quadratic effect on starch cook: too little water does not provide enough hydration of the starch granule and subsequently the kibble will have small cell structure and little expansion (3). On the other extreme, too much water decreases temperature in the extruder and therefore starch cook, acting as a plasticizer (32). When water is below what is required to hydrate the dry mix, the mechanical shear inside the extruder barrel can cause starch damage which will create a premature RVA peak in cold water, meaning that starch has been mechanically damaged. This phenomenon was present in the RVA profile of sample 16 (Fig 2). (3) demonstrated that 22% and 37% IBM were the extremes because neither resulted in kibble expansion or to have a good cell structure according to scanning electron micrographs. (33) reported that extruded wheat flour with the most RS was produced with excess water. At the other end of the spectrum, (34) reported that corn starch extruded at both high and low SS (600 and 300 rpm, respectively) at lower moisture content (17% water) yielded the most RS. In their study, the low water content was likely not enough to cook the starch, and the treatment with more water (at 22%) caused greater starch gelatinization. The treatment in the present study with high IBM, low SS and high PS yielded the greatest predicted RS content. The IBM at 36% likely helped to dissipate energy and lower starch cook. The process with higher moisture could also have created an environment to develop retrograded starch (RS type III) (33).

Extruder shaft speed (SS) is a controllable input that affects specific mechanical energy directly. Simultaneous to a higher SS and increased mechanical energy in the process the residence time decreases. This exposes the starch to a shorter cook time in the extruder barrel. Therefore, altering shaft speed can have mixed results. (35) reported that RS content of corn and mango were higher when extruded at a SS of 30 rpm as compared to 65 rpm. (33) did not find a difference in RS of wheat flour extruded at the same moisture contents and different shaft speeds. This is likely because they increased SS at 50 rpm increments, which was not a significant change to affect RS yield. In the present study, increasing the shaft speed in 400 rpm increments led to significant cross-product effects with particle size on RS yield.

Among methods used to determine the degree of starch gelatinization starch cook was the least consistent. This happened because even raw corn possesses pores and channels that facilitate enzyme adhesion and digestion (18). Thus, the starch cook method in corn does not account for only the portion that was gelatinized. A better method to estimate starch gelatinization in this study was RVA as it correlated highly with dough temperature and SME. The RVA provides a broader characterization of starch transformations during extrusion. With cold water raw starch does not swell and presents a low viscosity, while high molecular weight starch derivatives produced from chain scission during extrusion easily swell and increase viscosity (36). This was well illustrated in the RVA plots (Fig 2) where the treatment with the highest thermomechanical energy had a significant initial cold swelling and the less cooked sample had no cold swelling. As the RVA temperature increased to the gelatinization range of corn starch (mid 60ºC) (37) the sample that preserved some raw starch had an increase in relative viscosity, while the treatment that suffered more thermomechanical energy had little to no raw starch left to promote a hot viscosity peak. Samples 15 and 16 that were used as examples of RVA plots also had the highest and lowest IBM, respectively. The first sample had a much higher setback viscosity than the latter. According to (38), extruded puffs produced with high and low water contents had similar final viscosity as samples from the present study, which meant that less water led to more mechanical shear that dextrinized the starch, reducing final viscosity. Extrusion causes mechanical disruption of molecular bonds within the starch granule, resulting in loss of crystallinity and gelatinization (39,40). The RVA method to measure cooked and raw starch should be employed more frequently in pet foods, but it is important that the same protocol is used so that results can be comparable across studies.

Extrudate temperature is a result of thermomechanical energy inputs during the process. Mechanical energy is largely affected by viscosity, which is affected by water content. The driving force for bubble expansion in the kibble can be represented as a function of water vapor pressure inside the vapor bubble and viscosity (specific volume of extrudate = P_vs_/η, where P_vs_ is the water vapor pressure and η the molten viscosity) (41). In the present study both VEI and LEI had a high negative correlation with moisture content. This meant that the dough at 30% IBM likely had a lower viscosity compared to treatments produced at 35% IBM, which allowed for bubble expansion according to (41). Extrudates with high IBM won’t absorb as much mechanical energy and therefore won’t be as hot, so vapor pressure decreases (42), decreasing overall expansion. Moreover, a wetter extrudate takes a longer time to drop below glass transition temperature (Tg) and the shrinkage effect is greater (42).

## 5. Conclusion

This study was successful at describing the effect processing has on starch gelatinization and RS content in the kibble. Results suggested that a higher IBM, lower extruder shaft speed and larger particle size should contribute to the survival or development of RS during thermomechanical processing. Higher water content and lower extruder shaft speed lower the dough viscosity which directly affects SME. Resistant starch had large negative correlations with dough temperature and SME, which means that extrusion should target a low thermomechanical to increase RS yield. This strengthens the RS prediction model built in this study since SME is a function of multiple factors such as viscosity, and treatment inputs extruder shaft speed, water content and particle size. The physical method RVA to characterize starch gelatinization was preferred over the enzymatic starch cook as it had a strong correlation with thermomechanical parameters. The model created to predict RS can be used as a platform for future studies. Future work should focus on improving this model by using a wider range of corn particle sizes.

## Abbreviations

PC: extruder preconditioner
RS: resistant starch
RVA: rapid visco analyzer
SCFA: short-chain fatty acids
PS: particle size
OE: off the extruder
OD: off the drier
IBM: in-barrel moisture
SME: specific mechanical energy
SEI: sectional expansion index
VEI: volumetric expansion index
LEI: longitudinal expansion index
SS: extruder shaft speed.

## Author Contributions

Isabella Corsato Alvarenga (ICA), Christopher Waldy (CW) and Charles Gregory Aldrich (CGA) were responsible for study planning. ICA wrote the manuscript and conducted statistical analysis. CW did the experimental design and oversaw the extrusion processing. CGA and Lewis C. Keller (LCK) edited and provided intellectual contribution to the work.

## Acknowledgements

This work was supported by the Colgate-Palmolive Company. We would like to thank Dr. Dallas Johnson for statistical support.

